# Stepwise evolution of the developmental and symbiotic functions of DELLA in land plants

**DOI:** 10.64898/2026.03.02.709003

**Authors:** Katharina Melkonian, Tifenn Pellen, Katinka Bünger, Océane Thiercelin, Aurélie Le Ru, Mélanie K. Rich, Matheus E. Bianconi, Jean Keller, Pierre-Marc Delaux

## Abstract

DELLA proteins play diverse roles in development, plant-microbe interactions and stress responses, and are regulated by the hormone gibberellic acid (GA) in flowering plants. Here, we investigated the evolutionary conservation of functions of the single *DELLA* ortholog in the non-vascular model liverwort *Marchantia paleacea*, which lacks genes for perception and biosynthesis of bioactive GA. We found that GA-independent developmental phenotypes are conserved in *M. paleacea*, while neither *DELLA*, nor its close paralog *GRAS13* are involved in establishment of arbuscular mycorrhizal symbiosis. We propose that symbiosis-related functions of DELLA evolved and have been maintained in vascular plants because GA-dependent regulation offered an additional, systemic, control layer on symbiosis.

## Introduction

In flowering plants, DELLA proteins play multiple functions mediated by the plant hormone gibberellic acid (GA). DELLA are master growth-repressors and are subjected to degradation by the proteasome in presence of bioactive GA (Dill *et al*., 2004). High-yield dwarfed plants in wheat, maize and rice were found to have mutations in genes encoding for DELLA proteins and to be less responsive to GA (Peng *et al*., 1999; Hedden, 2003). The GA-induced degradation of DELLA proteins is mediated by the GA-receptor GID1 that was originally identified in rice (Ueguchi-Tanaka *et al*., 2005; Griffiths *et al*., 2007). DELLA proteins were shown to directly interact with Phytochrome-interacting factors (PIF) and thereby preventing the expression of PIF target genes (De Lucas *et al*., 2008; Feng *et al*., 2008). PIFs have been reported to play a key role in light-regulated plant development, for example mediated by phytochrome signalling and chlorophyll biosynthesis (Huq, 2002; Huq *et al*., 2004). Accordingly, DELLA proteins are also regulators of photomorphogenesis.

Beyond Arabidopsis, DELLA, GID1 and bioactive GAs have been detected in all other vascular plant lineages (Hirano *et al*., 2007; Yasumura *et al*., 2007). By contrast, both GID1 and GA bioactive in vascular plants are absent from bryophytes (Rensing *et al*., 2008; Bowman *et al*., 2017; Rich *et al*., 2021). Yet, knocking-out *DELLA* in the moss *Physcomitrium patens* or over-expression of DELLA in the liverwort *Marchantia polymorpha* lead to developmental defects (Phokas *et al*., 2023; Hernández-García *et al*., 2021). *P. patens* mutants are impaired in sporophyte development and display an increased spore germination rate, while no clear role in light signaling has been identified. *M. polymorpha DELLA^OE^* lines were found to be smaller than wildtype plants and overexpression of *DELLA* caused a loss of gemma dormancy, which led to early germination of gemmae inside gemma cups. In addition, *M. polymorpha DELLA^OE^* lines are delayed in sexual reproduction.

In flowering plants, DELLA was also identified as a positive regulator of symbiosis establishment with diverse microorganisms, such as nitrogen-fixing rhizobia or arbuscular mycorrhizal (AM) fungi (Floss *et al*., 2013; Yu *et al*., 2014; Jin *et al*., 2016). For instance, in the legume *Medicago truncatula*, *della* mutants show decreased colonization by AM fungi, and a lack of arbuscule formation (Floss *et al*., 2013; Jin *et al*., 2016). In rice, the effect is even more pronounced, with the almost total absence of colonization (Yu *et al*., 2014). Protein-protein interaction studies proposed that DELLA is part of a nuclear-localized complex with components of the common symbiosis pathway essential for integrating chemical signals produced by symbiotic microorganisms (Jin *et al*., 2016). Because the two model non-vascular plants *P. patens* and *M. polymorpha* have lost the ability to engage in the ancestral mutualistic symbiosis with AM fungi (Radhakrishnan *et al*., 2020), the evolutionary conservation of the symbiotic function of DELLA has never been investigated.

The liverwort *Marchantia paleacea* has become model of choice to study the conservation of the AM symbiosis across land plants by comparison with vascular plants. So far, all the tested symbiotic mechanism were found conserved across land plants, including the perception of fungal-derived chemical signals (Teyssier *et al*., 2025; Tan *et al*., 2025), the integration of these signals by the common symbiosis pathway (Lam *et al*., 2024; Vernie *et al*., 2025; Rich *et al*., 2025), arbuscule formation (Sgroi *et al*., 2024), and the transfer of lipids to the fungal symbiont (Rich *et al*., 2021).

In this study, we generated CRISPR/Cas9-mediated *M. paleacea della* mutants, as well as double mutants (*Mpadella/gras13)* with its closely related *GRAS13* paralog. Phenotyping revealed that the origins of the developmental and symbiotic functions of DELLA were uncoupled during the evolution of land plants.

## Results and Discussion

### The single *DELLA* gene plays a developmental role in *M. paleacea*

To understand the function of DELLA in *M. paleacea,* we first generated *MpaDELLA_pro_:GUS* lines and monitored the DELLA_pro_ activity. In mature thalli, blue staining from the GUS-reporter was mainly observed in the photosynthetic layer, with a strong focus in assimilatory filaments at air pores and in rhizoids (Figure 1f). Next, we aimed at generating *M. paleacea della* mutants. Previous attempts in *M. polymorpha* did not allow generating viable *della* mutants, limiting the functional characterization to lines over-expressing DELLA (Hernández-García *et al*., 2021). Using CRISPR/Cas9, we managed to generate five *della* mutants in *M. paleacea* (Figure S1a, d). Mutations ranged from single base substitutions to 62 base deletions (Figure S1a, d). Because the growth of *M. paleacea* is not homogenous *in vitro* we conducted the developmental phenotyping in more realistic conditions, similar to the promoter GUS conditions (Material and Methods). We monitored the growth of *Mpadella* mutants over time, which revealed a growth retardation of *della* mutants compared to control plants (Figure 1g). At 57-days old, all five *della* mutant alleles were significantly smaller than control plants (Figure 1a, b). We quantified the number of gemma cups at 57-days old and found that four out of five *della* mutant alleles produced significantly less gemma cups than control plants (Figure 1d). In addition, three out of five *della* mutant alleles appeared slightly less green than control plants based on image quantification at 57-days old (Figure 1c), which prompted us to determine the relative Chlorophyll a and Chlorophyll b absorbance in these plants (Figure 1e). We found that both Chlorophyll a and Chlorophyll b absorbance were significantly lower in all five mutant alleles compared to control plants (Figure 1e). Over-expression experiments in *M. polymorpha* suggested that DELLA might play a role integrating growth and light responses (Hernández-García *et al*., 2021). To further test this hypothesis, empty-vector control lines and *della* mutants were grown under medium (75 µmol photons m^-2^ s^-1^) and high (100 µmol photons m^-2^ s^-1^) light regimes. In control lines, this treatment resulted in an overall similar growth speed compared to normal light conditions for medium light regime and a slightly increased growth speed at high light regime (Figure S2g, Figure S3g). In contrast, in *della* mutant plants, higher light intensities led to a stronger, but highly non-homogenous increase of growth speed, with asymmetric growth patterns of thalli (Figure S2a,b, Figure S3a,b). This effect was stronger at medium light regime, leading to an overall lower difference in size at 57-days old of the *della* mutants compared to control plants (Figure S2b). A similar effect was observed for green coloration based on image quantification (Figure S2c, Figure S3c) as well as gemma cup quantification (Figure S2d, Figure S3d). In the case of relative Chlorophyll a and Chlorophyll b absorption, less differences between *della* mutant plants and control plants were observed at high light regime (Figure S2e Figure S3e). The expression pattern of *DELLA,* as per promoter-GUS reporter lines, did not drastically change between different light regimes, but appeared more focused at high light regime (Figure 1f, Figure S2f, Figure S3f).

**Figure 1:**
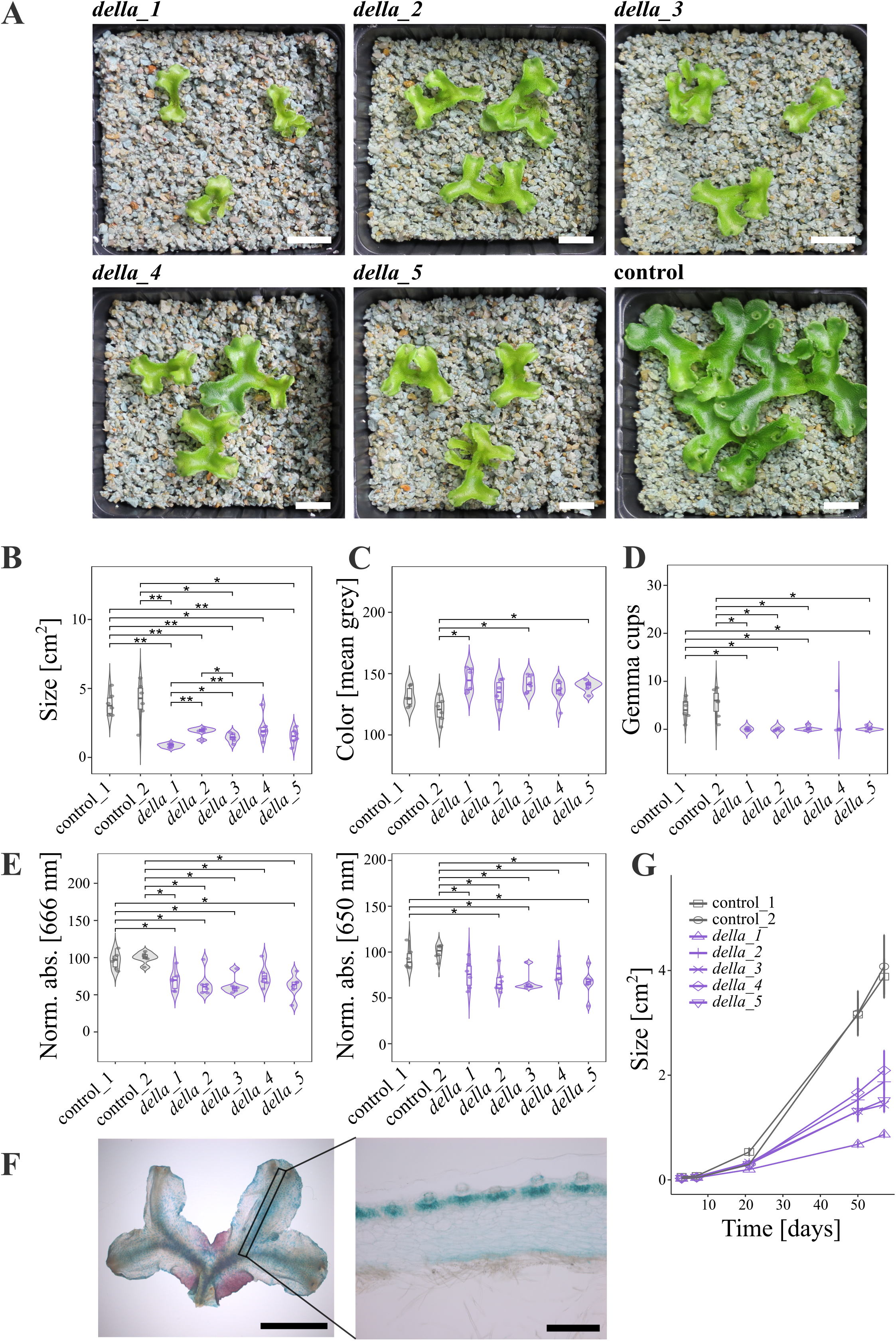
Mpa*della* mutants are impaired in development. **A.** Representative overview pictures of Mpa*della* and control plants at 57-days old, grown at 50 µmol photons m^-2^ s^-1^. Scalebar: 1cm. **B.** Thallus size at 57-days old (n=6). **C.** Mean grey value of thallus-area (n=6). **D.** Gemma cup number per genotype (n=6). **E.** Relative chlorophyll a and chlorophyll b absorption. Combined results of two independent experiments are shown (n=6). For all measures in **B.-E.**, statistics were performed using pairwise Wilcoxon rank sum test and p-values were corrected using the fdr method (p ≤ 0.05 ∼ “*”; p ≤ 0.01 ∼ “**”; p ≤ 0.001 ∼ “***”; p ≤ 0.0001 ∼ “****”). **F.** Promoter-GUS reporter expression in proMpa*DELLA*:*GUS* lines at 57-days old. Left: Overview, scalebar: 1 cm. Right: Longitudinal section, scalebar: 0.5 mm. **G.** Thallus size quantification over the course of 57 days. Data are mean +/- SE (n=6). Two independent repetitions of the experiment were performed.

We propose that DELLA is a positive regulator of growth and gemma cup formation in *M. paleacea*, which are regulated in a light-dependent manner to some extend and might be linked to PIF signalling (Hernández-García *et al*., 2021) and chlorophyll biosynthesis.

### *MpaDELLA* is not required for the arbuscular mycorrhizal symbiosis

In flowering plants, mutations in *della* lead to very significant symbiotic defects, including abolished or reduced colonization, and the lack of arbuscule formation (Floss *et al*., 2013; Yu *et al*., 2014). We first analyzed the *MpaDELLApro:GUS* lines in *Rhizophagus*-inoculated plants. Compared to the mock-inoculated condition, an additional faint signal was observed in the colonized area of the thallus (Figure 2a). To investigate the symbiotic function of *DELLA* in *M. paleacea* during symbiosis, empty-vector control plants and the five *della* mutants were inoculated with the arbuscular mycorrhizal fungus *Rhizophagus irregularis*. Six weeks after inoculation, the thalli were harvested and AM colonization assessed. We initially scored the percentage of infected plants based on the presence or absence of an AMS-induced purple pigment as a proxy for fungal colonization (Vernié *et al*., 2025). We found that one out of five mutant alleles displayed reduced infection by *Rhizophagus irregularis*, while another allele displayed significantly increased infection compared to control_1 plants (Figure 2c). We conclude that these observations are rather caused by experimental variation or differences in loci of T-DNA insertion than based on *DELLA* functions. Next, we quantified the progression of fungal colonization. We found that fungal colonization was significantly less progressed in four out of five *della* mutant alleles compared to control plants (Figure 2d). Given the observed developmental defects of *della* mutant alleles (Figure 1a, S2a, S3a, 2b), these findings should be interpreted with caution and might be the result of a growth retardation or reduced photosynthetic capacity of the *della* mutants compared to control plants. Finally, we analyzed the development of the fungal structures in the thalli. We found that arbuscules were present in all five *della* mutant alleles with no morphological impairments (Figure S4b), which is in sharp contrast to the findings described in model flowering plants (Floss *et al*., 2013; Jin *et al*., 2016).

**Figure 2:**
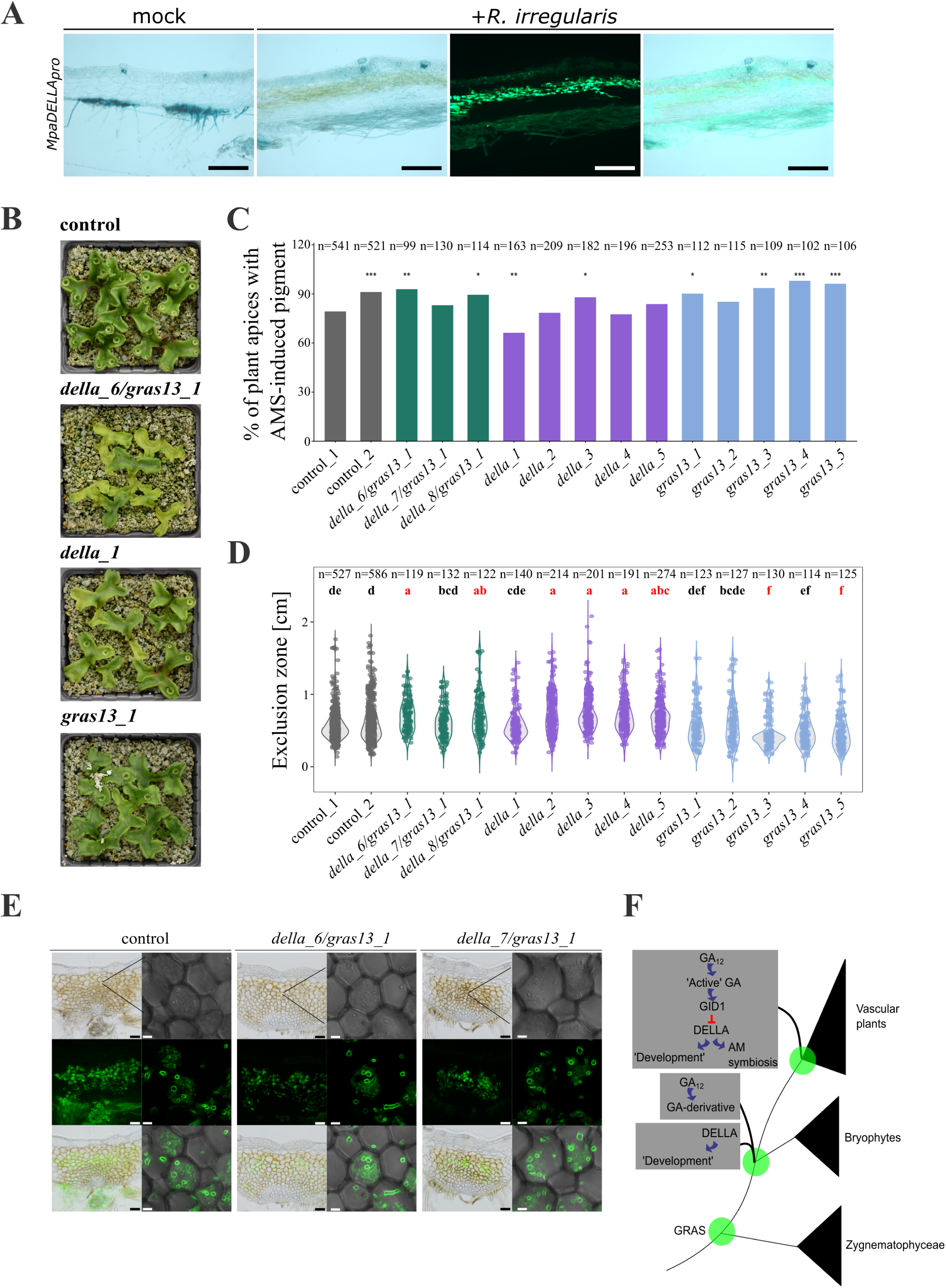
Mpa*DELLA* is not essential for arbuscular mycorrhizal symbiosis in *M. paleacea*. **A.** Promoter activity of the *Marchantia paleacea DELLA* monitored using Mpa*DELLA*pro:*GUS* at 6 weeks post inoculation. From left to right, longitudinal section of mock inoculated (bright field), *R.irregularis*-inoculated (bright field), *R.irregularis*-inoculated (WGA-Alexafluor 488), *R.irregularis*-inoculated (merged) plants. **B.** Representative images of control plants transformed with an empty vector, Mpa*della/gras13*, Mpa*della* and Mpa*gras13* plants inoculated with *R. irregularis* at 6 weeks post inoculation. **C.** Scoring of plant apexes with AMS-induced pigment at 6 weeks post inoculation with *R. irregularis* in control plants transformed with an empty vector and Mpa*della/gras13*, Mpa*della* and Mpa*gras13* mutants. Statistics were performed using pairwise Chi^2^ test with control_1 plants as reference group and p-values were corrected using the Benjamini Hochberg method. Combined results of 6 independent experiments are shown. **D.** Progression of colonization measured by exclusion zone at 6 weeks post inoculation with *R. irregularis* in control plants transformed with an empty vector and Mpa*della/gras13*, Mpa*della* and Mpa*gras13* mutants. Combined results of 6 independent experiments are shown. Statistics were performed using One Way Anova and Tukey HSD test. Red letters indicate statistically significant differences from control plants. **E.** Representative images of control plants transformed with an empty vector and Mpa*della/gras13* double mutants inoculated with *R. irregularis* at 6 weeks post inoculation. Green channel: WGA-Alexafluor 488 staining of fungal structures. Scale bar 100 μM or 10 μM. **F.** Proposed model for the step-wise evolution of DELLA functions in the green lineage. GRAS were gained in *Zygnematophyceae* via HGT (Cheng *et al,* 2019). In bryophytes DELLA functions independently of GA and it’s precursors in development. GA-dependent regulation of DELLA evolved in vascular plants and was co-opted as an additional layer of control during arbuscular mycorrhizal symbiosis.

We conclude from the analysis of the five *M. paleacea della* mutants that their altered growth leads to a quantitative decrease in overall colonization, yet the formation of arbuscule is not impaired.

### *MpaDELLA* and its paralog *MpaGRAS13* are not redundant for AM symbiosis

The lack of an AM symbiosis defect similar to angiosperms in the *M. paleacea della* mutants might either indicate that symbiosis signalling differs between angiosperms and bryophytes, or be the result of functional redundancy. Our phylogenetic analysis conducted on *DELLA* homologs confirmed the existence of a liverwort-specific *DELLA* paralog (Hernández-García et al. 2019), namely Mpa*GRAS13*, which likely evolved from a duplication in bryophytes followed by subsequent losses in mosses and hornworts (Figure S5). To determine whether *M. paleacea GRAS13* could act redundantly with *DELLA*, we generated single *gras13* and *della/gras13* double mutants (Figure S1b, c, e, f), and phenotyped them in presence of AM fungi. No strong developmental phenotypes were observed in the case of *gras13* single mutants, while the *della/gras13* double mutants displayed similar developmental phenotypes as observed for *della* single mutants, but to a much stronger extend (Figure 2b, S4a). In particular, *della/gras13* double mutants showed a strong asymmetric growth pattern and a complete absence of gemma cups in all three mutant alleles (Figure S4a). When scoring AM symbiosis using the AM symbiosis-induced pigment, we found that four out of five *gras13* mutant alleles and two out of three *della/gras13* double mutant alleles displayed a significantly increased percentage of infected plants compared to one of the two control lines (control_1) (Figure 2c). However, when compared to the second control line (control_2), this effect was in the range of biological variation for this type of experiment. Next, we quantified the progression of fungal colonization in *gras13* single and *della/gras13* double mutants. We found that fungal colonization was significantly increased in two out of five *gras13* single mutant alleles and significantly delayed in two out of three *della/gras13* double mutant alleles compared to control plants (Figure 2d). Intracellular arbuscules were present in all *gras13* single and all *della/gras13* double mutant alleles, with no morphological defects compared to control plants (Figure 2e, S4b).

We conclude from the analysis of the *M. paleacea della/gras13 double* mutants that the developmental phenotypes observed leads to a quantitative decrease in overall colonization, yet the formation of arbuscule is not impaired.

### A model for the evolution of DELLA function in land plants

Biochemical surveys coupled with phylogenetic analyses of the Gibberelic Acid (GA) biosynthesis and signalling pathways revealed that the evolution of DELLA predated the production of canonical GAs (Sun *et al*., 2023). Indeed, while DELLA and the upstream part of the biosynthesis, down to *ent-kaurenoic acid* and the intermediate GA_12_ (Sun *et al*., 2023), are present in non-vascular plants, the GA receptor GID1 and GA sensing is specific to the vascular plants (Hirano *et al*., 2007). This points to the evolution of a GA-GID1-DELLA module at the base of vascular plants (Hirano *et al*., 2007; Sun *et al*., 2023).

The five *M. paleacea della* mutants generated in this study confirmed the developmental role of DELLA in *Marchantia* previously suggested by over-expression (Hernández-García *et al*., 2021). Together with the developmental defect of the *P. patens della* mutants (Phokas *et al*., 2023), this indicates that DELLA plays a GA-independent developmental role in non-vascular plants.

By contrast with the developmental function, the lack of a strong AM symbiosis defect in the *M. paleacea della* and *della/gras13* mutants demonstrates that DELLA is not involved in symbiosis in non-vascular plants. This result is intriguing on its own. So far, all the symbiotic genes identified in angiosperms and tested for their symbiotic functions revealed a conserved role in *M. paleacea* (Sgroi *et al*., 2024; Lam *et al*., 2024; Teyssier *et al*., 2025; Vernie *et al*., 2025; Rich *et al*., 2025), including components acting together with DELLA in angiosperms such as CYCLOPS and CCaMK (Vernié *et al*. 2025).

It can be proposed that during the evolution of vascular plants, GA_12_-modifying enzymes emerged, leading to the biosynthesis of canonical GAs. The existing DELLA-regulated module was co-opted by these GAs via the gain of a functional GID1 receptor (Hirano *et al*., 2007; Yasumura *et al*., 2007; Yoshida *et al*., 2018). Interestingly, the GA intermediate identified in *M. polymorpha,* GA_12_, is the form transported via the vasculature in angiosperms (Da Fonseca *et al*., 2015). Beyond symbiosis, systemic regulations allowing for long-distance communication across organs and tissue was one of the key innovations for the vascular plants to evolved (Wheeldon and Bennett, 2021). We propose the hypothesis that DELLA has evolved and been maintained in the vascular plant symbiotic signalling pathway not directly because of its positive role, but rather because its degradation by GA brings an additional, systemic, control layer on symbiosis.

## Acknowledgements

Research performed at LRSV was also supported by the “Laboratoires d’ Excellence (LABEX)’ TULIP (ANR-10-LABX-41) and the “École Universitaire de Recherche (EUR)” TULIP-GS (ANR-18-EURE-0019). This work was supported by the German Research Foundation (DFG Walter Benjamin fellowship project number 536856410 to KM); the Horizon Europe programme (MSCA-PF grant 101105838 ‘SYMBIOLOSS’ to MEB); and the European Research Council (ERC) under the European Union’s Horizon 2020 research and innovation programme (grant agreement no.101001675-ORIGINS) to P.-M.D.

## Author contributions

KM, KB, OT conducted and analyzed the experiments. TP, KM, MKR generated the genetic resources. JK, MEB conducted and analyzed phylogenies. PMD, KM conceived and coordinated the project. PMD, KM wrote the manuscript.

## Data availability

All data are provided in the manuscript and supplementary materials.

## Supporting information

**Table S1. Plasmids and guideRNA sequences.**

**Table S2. Supplementary data for Mpa*della* experiments at different light regimes.**

**Table S3. List of land plant species genomes used for the phylogenetic analyses.**

## Material and methods

### Stable transformation of *Marchantia paleacea*

Genetic constructs for CRISPR-Cas9 mediated targeted mutagenesis and promoter-reporter assays were generated using the Golden Gate molecular cloning system as previously described (Engler and Marillonnet, 2014; Patron *et al*., 2015; Rich *et al*., 2021). A list of guideRNA sequences, as well as Level 0, 1, and 2 constructs is provided in Supplementary Table S1. *M. paleacea* wildtype plants were grown on Gamborgs B5 half strength medium (G5768, Sigma, France) pH 5.7, 1.4% bacteriological agar (1330, Euromedex, France) for 4 to 5 weeks in a walk-in growth chamber at 22 °C under 16 h light/8 h dark cycle. For each transformation, 15-25 gemmalings were blended in a sterile 250 ml stainless steel bowl (Waring, USA) in 10 ml of 0M51C medium (KNO_3_ 2 g/L, NH_4_NO_3_ 0.4 g/L, MgSO_4_ 7H_2_O 0.37 g/L, CaCl_2_ 2H_2_O 0.3 g/L, KH_2_PO_4_ 0.275 g/L, L-glutamine 0.3 g/L, casamino-acids 1 g/L, Na_2_MoO_4_ 2H_2_O 0.25 mg/L, CuSO_4_ 5H_2_O 0.025 mg/L, CoCl_2_ 6H_2_O 0.025 mg/L, ZnSO_4_ 7H_2_O 2 mg/L, MnSO_4_ H_2_O 10 mg/L, H_3_BO_3_ 3 mg/L, KI 0.75 mg/L, EDTA ferric sodium 36.7 mg/L, myo-inositol 100 mg/L, nicotinic acid 1 mg/L, pyridoxine HCl 1 mg/L, thiamine HCl 10 mg/L). The plant fragments were cultivated in 250ml erlenmeyer flasks containing 15 ml 0M51C for 3 days with shaking (200rpm) at 22 °C under 16 h light/8 h dark cycle. The plant fragments were subsequently inoculated with 100 µL *Agrobacterium tumefaciens* GV3101 harbouring the respective plasmids from a saturated liquid culture and supplemented with acetosyringone to a final concentration of 100 µM. The plant fragments were co-cultivated with *Agrobacteria* for 3 days with shaking (200rpm) at 22 °C under 16 h light/8 h dark cycle, washed three times with water and plated on Gamborgs B5 half strength solid medium containing 200 mg/L amoxycilin (Levmentin, Laboratoires Delbert, FR) and 10 mg/L Hygromycin (Duchefa Biochimie, France). Transformants were selected after 3-5 weeks cultivation at 22 °C under 16 h light/8 h dark cycle.

### Phenotyping of Mpa*della* mutants at different light regimes

Three *Marchantia paleacea* plants were grown in two pots per genotype on zeolite substrate (50% fraction 1.0 to 2.5 mm, 50% fraction 0.5 to 1.0-mm, Symbiom) at 22-24 °C in a walk-in growth chamber under long-day conditions (16hr light/8hr dark) at different light intensities (low: 50 µmol photons m-2 s-1; medium: 75 µmol photons m-2 s-1; high: 100 µmol photons m-2 s-1). The plants were watered once per week with Long Ashton solution containing 15 µM phosphate and plant growth was monitored over the course of 57 days. Two independent repetitions of the experiment were performed. Plant size and mean grey values were quantified using ImageJ. Gemma cup number was quantified on day 57 of the experiment. For Chlorophyll absorbance determination, three plants per genotype and condition were harvested and weighed on day 57 of the experiment. Chlorophyll was sequentially extracted from the plant tissues in 2 ml DMSO at 65 °C for 4 hrs in the dark. Chlorophyll absorbance in the supernatant was determined using a photometer (P3 UVisco, LC Instrument) at a wavelength of 666 nm for Chlorophyll a and 650 nm for Chlorophyll b, respectively. Absorbance values were normalized according to plant fresh weight per sample and relative values were calculated through division by the mean absorbance of control plants per sample. For all measures, statistics were performed using pairwise Wilcoxon rank sum test and p-values were corrected using the fdr method (p ≤ 0.05 ∼ “*”; p ≤ 0.01 ∼ “**”; p ≤ 0.001 ∼ “***”; p ≤ 0.0001 ∼ “****”). Raw counts and calculated values are provided in Supplementary Table S2. All data were plotted using R Studio (version 2025.05.0).

### *Marchantia paleacea* mycorrhization tests and promoter-GUS reporter assays

*Marchantia paleacea* plants were grown in pots on zeolite substrate (50% fraction 1.0 to 2.5 mm, 50% fraction 0.5 to 1.0-mm, Symbiom) at 22-24 °C in a walk-in growth chamber under long-day conditions (16hr light/8hr dark) at 65 µmol photons m^-2^ s^-1^ for two weeks. Plants were watered once per week with Long Ashton solution containing 15 µM phosphate. Five plants per pot were then mock-inoculated or inoculated with approximately 1000 sterile spores of *Rhizophagus irregularis* DAOM 197198 provided by Agronutrition (Labège, France, https://www.agronutrition.com/en/contact-us/). At 6 weeks post inoculation, the plants were harvested. For promoter-GUS reporter assays, the plants were vacuum-infiltrated with GUS buffer (phosphate buffer (0.1 M), EDTA (5 mM), K_3_Fe(CN)_6_ (0.5 mM), K_4_Fe(CN)_6_ (0.5 mM), X-Gluc (0.25 mg/ml, Euromedex, France), and H_2_O) for 20 min at 200 mbar directly after harvesting and then incubated at 37 °C overnight in the dark. The GUS buffer was removed and the plants were washed once with water. All plants were then submerged in 100% ethanol to extract chlorophyll until transparency was achieved and then stored in aqueous solution containing EDTA (0.5 mM) until further processing. Cleared thalli were scanned and scored as previously described (Rich *et al*., 2021), based on presence or absence of a mycorrhization-induced purple pigment. Statistics were perfomed using Chi^2^ test and p-values were corrected using the Benjamini Hochberg method. Progression of colonization was determined by measuring the exclusion zone, as previously described (Rich *et al*., 2021). Statistics were performed using One Way Anova and Tukey HSD test. All data were plotted using R Studio (version 2025.05.0). The thalli were subsequently analyzed by microscopic imaging to confirm the presence of intracellular fungal structures.

### Microscopic imaging

Brightfield images of whole thalli in promoter-GUS reporter assays were acquired using an Zeiss Axiozoom V16 equipped with objective 0.5x. To visualize arbuscules and tissue-specific expression patterns, thalli were embedded in 7% agarose and 100 μM transversal or longitudinal sections were prepared using a Leica vt1000s vibratome. The sections were incubated overnight in 12% KOH at room temperature and rinsed 3 times with water, followed by an incubation in 1μg/ml WGA-Alexa Fluor 488 (Invitrogen, France) in PBS buffer overnight at 4 °C in the dark. The sections were then imaged using a Zeiss Axiozoom V16 equipped with objective 0.5x and GFP like filter set (Zeiss 38 He filter set, ex:470/40, em: 525/50) to visualize WGA or using a Nikon Ti Eclipse inverted microscope equipped with DS Ri2 camera and motorized XY stage. Large images of global sections were generated using the NISAR 4.3 scan large image module that allow multifield acquisition and images stitching. These images were acquired with 10×/0.3 dry objective (0.73 pixel size) in brightfield and in fluorescence for WGA-Alexa 488 staining using a GFP band pass filter set (ex: 472/30 nm, em:520/35 nm). Close-up images were acquired using a Leica SP8 TCSPC confocal microscope and LAS X software with a 25× water immersion objective (Fluotar VISIR 25×/0.95 WATER) at zoom 5× (0.182 μm pixel size). WGA-Alexa 488 staining of fungal structures was excited with the 488 nm laser line and fluorescence was recovered between 500 nm and 550 nm. Brightfield images were also performed and merged. Images were processed with ImageJ.

### Phylogenetic analyses

We used BLAST v2.15.0 (Camacho *et al*., 2009) to search for putative homologs of DELLA proteins in the genomes of 26 land plant species, including 14 vascular plants and 12 bryophytes (Supplementary Table S3), using the protein sequences of *Marchantia paleacea* as queries (*e-value* = 10^-9^). Sequences were aligned using MAFFT v7.505 (Katoh and Standley, 2013) with the automatic mode (*--auto*, *--maxiterate* 20), and subsequently trimmed using trimAl v1.4.1 (Capella-Gutiérrez *et al*., 2009) to remove alignment columns with more than 80% of gaps (*-gt* = 0.2). A phylogenetic tree was then estimated using FastTree v2.1.11 (Price *et al*., 2010) with the gamma model (*-gamma*). This resulting tree was inspected to identify the ortholog group containing the query sequences, which was extracted and realigned with the option *--maxiterate* set to 1000 to increase alignment accuracy, and trimmed to remove columns with more than 75% of gaps (*-gt* 0.25). A maximum-likelihood tree was then estimated with IQ-TREE v2.2.2.6 (Minh *et al*., 2020) using the standard model selection (*-m TEST*), with branch support assessed using the SH-like approximate likelihood test and ultrafast bootstrap, both with 1000 replicates.

---

**Fig S1.**
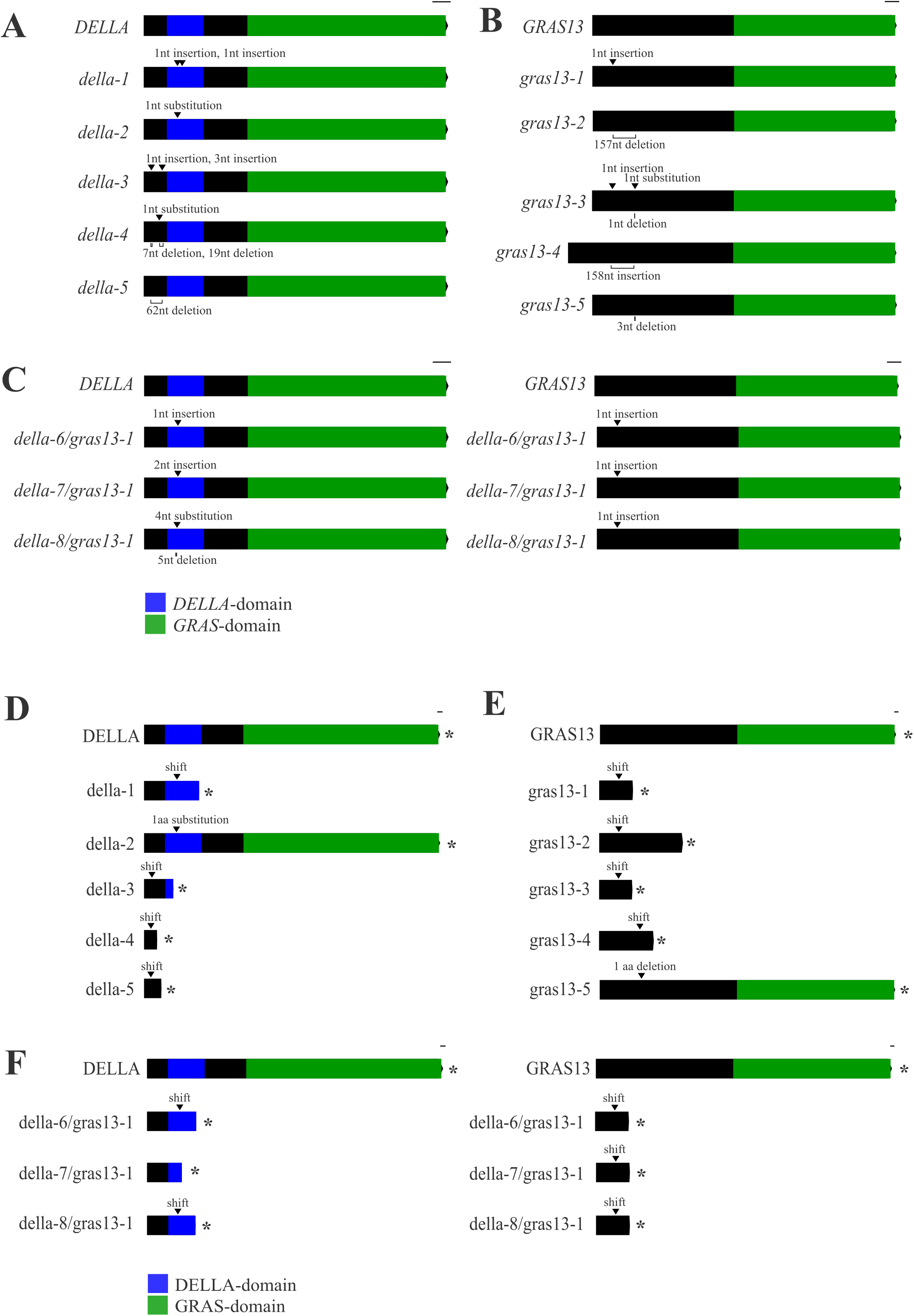
Gene and protein models of Mpa*della*, Mpa*gras13* and Mpa*della/gras13* mutants. **A.**, **B.** Schematic overview of mRNA structure of **A.** wildtype Mpa*DELLA* and mutated versions and **B.** wildtype Mpa*GRAS13* and mutated versions. Black arrows indicate the sites of mutations. **C.** Schematic overview of mRNA structure of wildtype Mpa*DELLA*, Mpa*GRAS13* and Mpa*DELLA/GRAS13* double mutants. Black arrows indicate the sites of mutations. **A.-C.** Scalebar: 100 basepairs. **D.**, **E.** Schematic overview of predicted proteins of **D.** wildtype MpaDELLA and mutated versions and **E.** wildtype MpaGRAS13 and mutated versions. **F.** Schematic overview of predicted proteins of wildtype MpaDELLA, MpaGRAS13 and MpaDELLA/GRAS13 double mutants. **D.-F.** Black arrows indicate sites of mutations and stars indicate predicted stop codons; Scalebar: 10 aminoacids. Gene and protein models were generated with http://wormweb.org/exonintron.

**Fig S2.**
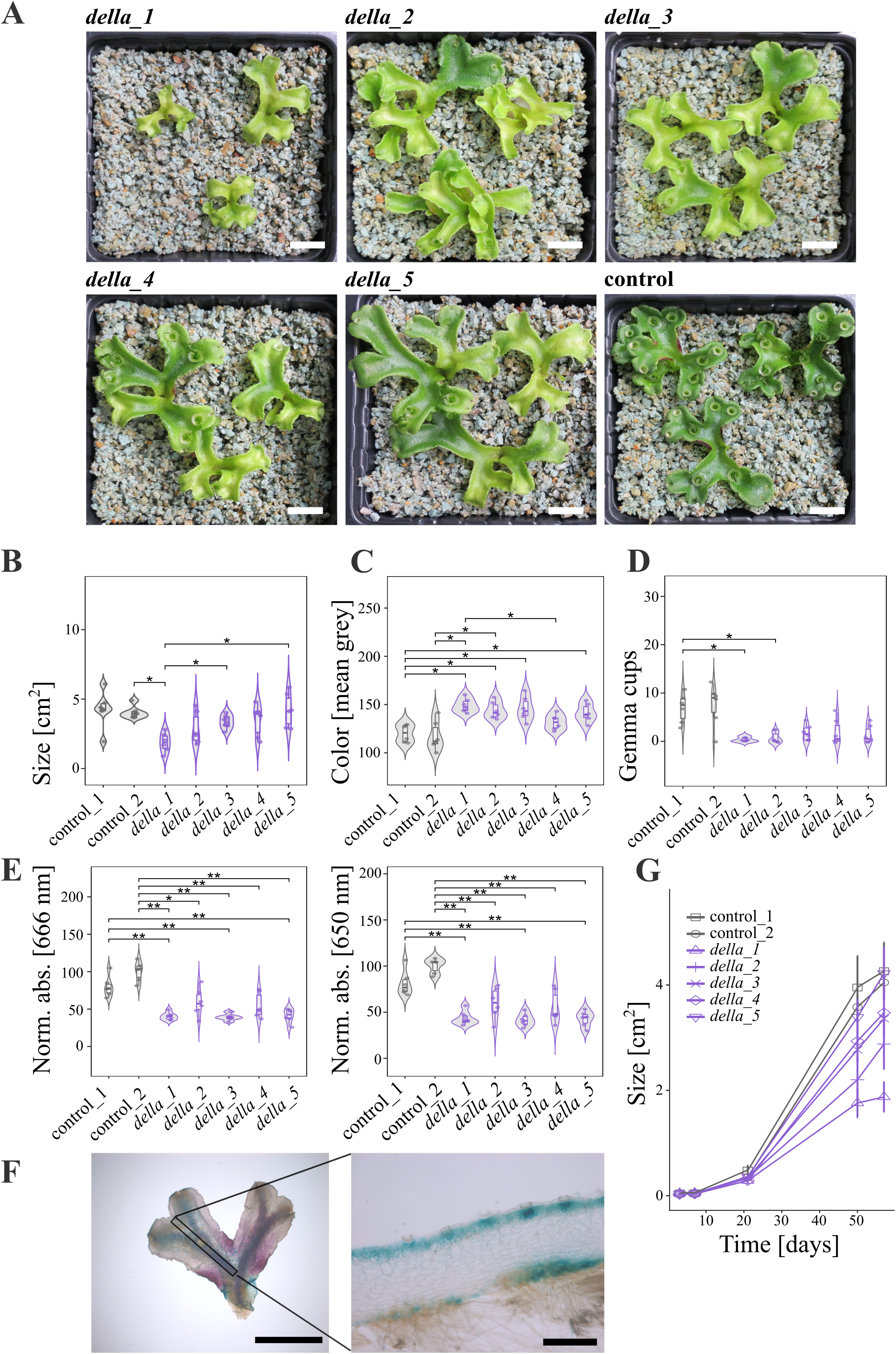
Phenotypic characterization of Mpa*della* mutants at medium light regime. **A.** Representative overview pictures of Mpa*della* and control plants at 57-days old, grown at 75 µmol photons m^-2^ s^-1^. Scalebar: 1cm. **B.** Thallus size at 57-days old (n=6). **C.** Mean grey value of thallus-area (n=6). **D.** Gemma cup number per genotype (n=6). **E.** Relative chlorophyll a and chlorophyll b absorption. Combined results of two independent experiments are shown (n=6). For all measures in **B.-E.**, statistics were performed using pairwise Wilcoxon rank sum test and p-values were corrected using the fdr method (p ≤ 0.05 ∼ “*”; p ≤ 0.01 ∼ “**”; p ≤ 0.001 ∼ “***”; p ≤ 0.0001 ∼ “****”). **F.** Promoter-GUS reporter expression in proMpa*DELLA*:*GUS* lines at 57-days old. Left: Overview, scalebar: 1 cm. Right: Longitudinal section, scalebar: 0.5 mm. **G.** Thallus size quantification over the course of 57 days. Data are mean +/- SE (n=6). Two independent repetitions of the experiment were performed.

**Fig S3.**
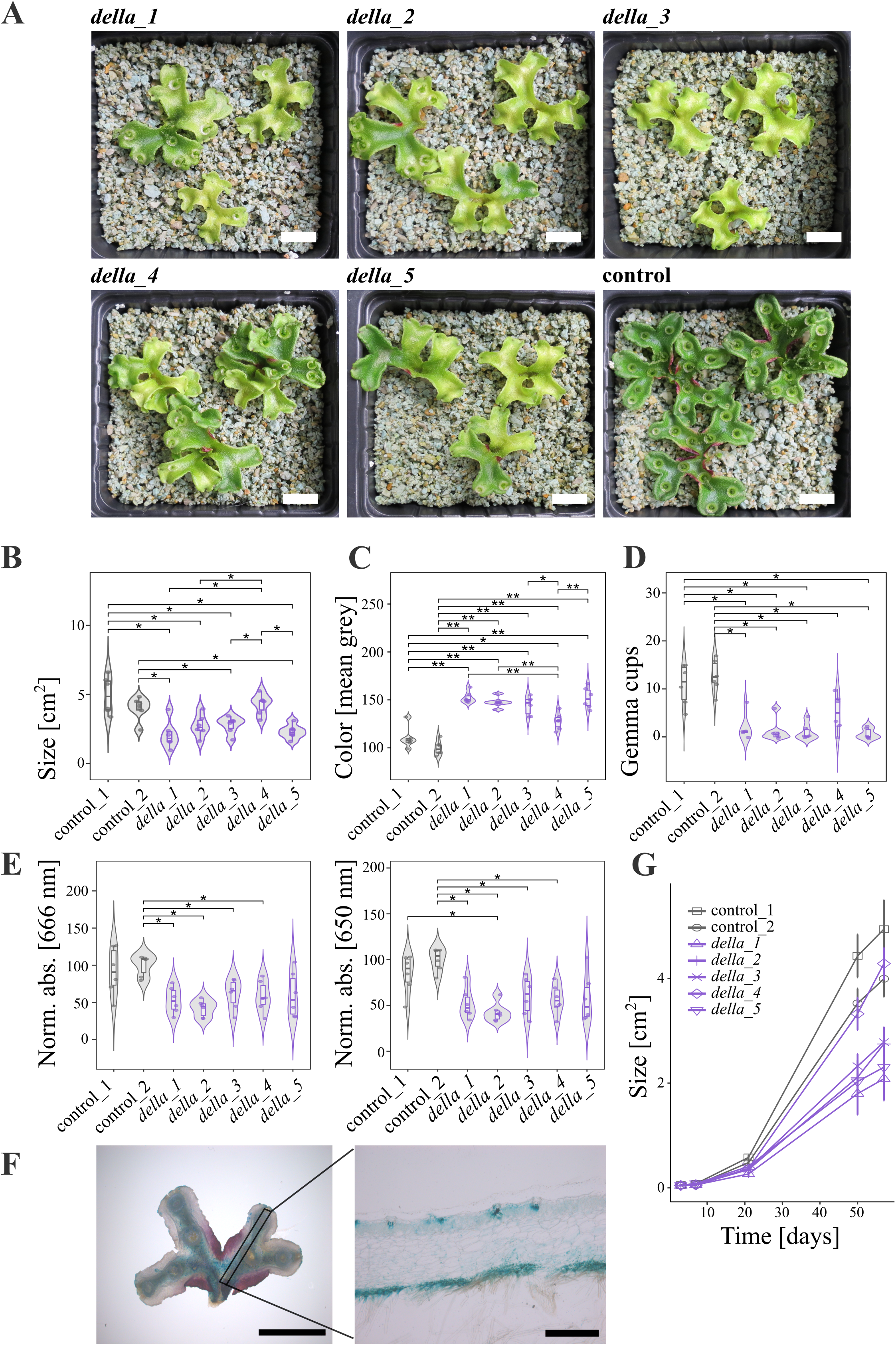
Phenotypic characterization of Mpa*della* mutants at high light regime. **A.** Representative overview pictures of Mpa*della* and control plants at 57-days old, grown at 100 µmol photons m^-2^ s^-1^. Scalebar: 1cm. **B.** Thallus size at 57-days old (n=6). **C.** Mean grey value of thallus-area (n=6). **D.** Gemma cup number per genotype (n=6). **E.** Relative chlorophyll a and chlorophyll b absorption. Combined results of two independent experiments are shown (n=6). For all measures in **B.-E.**, statistics were performed using pairwise Wilcoxon rank sum test and p-values were corrected using the fdr method (p ≤ 0.05 ∼ “*”; p ≤ 0.01 ∼ “**”; p ≤ 0.001 ∼ “***”; p ≤ 0.0001 ∼ “****”). **F.** Promoter-GUS reporter expression in proMpa*DELLA*:*GUS* lines at 57-days old. Left: Overview, scalebar: 1 mm. Right: Longitudinal section, scalebar: 0.5 mm. **G.** Thallus size quantification over the course of 57 days. Data are mean +/- SE (n=6). Two independent repetitions of the experiment were performed.

**Fig S4.**
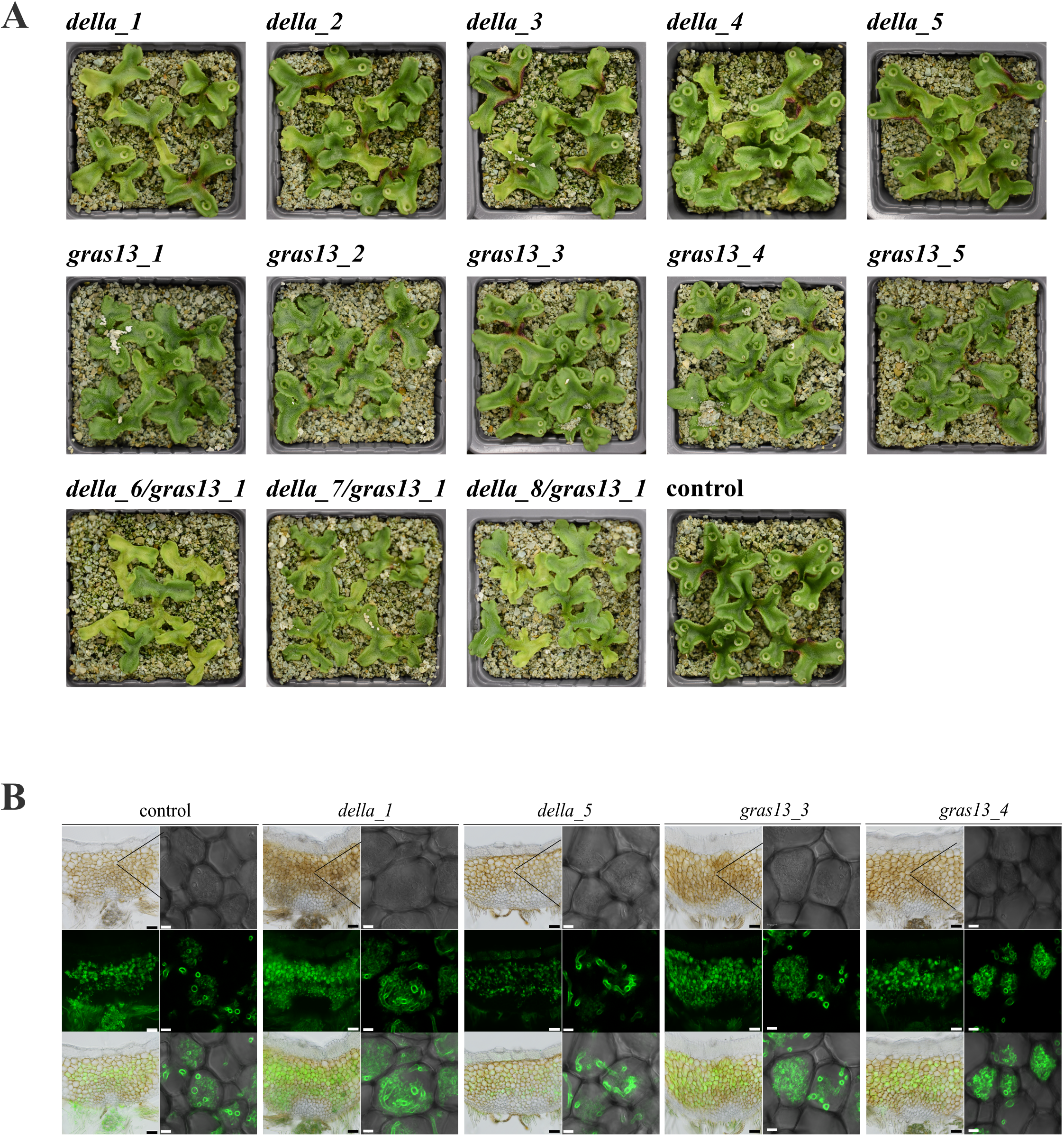
Macroscopic and microscopic phenotypes of Mpa*della*, Mpa*gras13* and Mpa*della/gras13* mutants. **A.** Representative pictures of control plants transformed with an empty vector, Mpa*della/gras13*, Mpa*della* and Mpa*gras13* plants inoculated with *R. irregularis* at 6 weeks post inoculation. **B.** Representative microscopic images of control plants transformed with an empty vector and Mpa*della/gras13* double mutants inoculated with *R. irregularis* at 6 weeks post inoculation. Green channel: WGA-Alexafluor 488 staining of fungal structures. Scale bar 100 μM or 10 μM.

**Fig S5.**
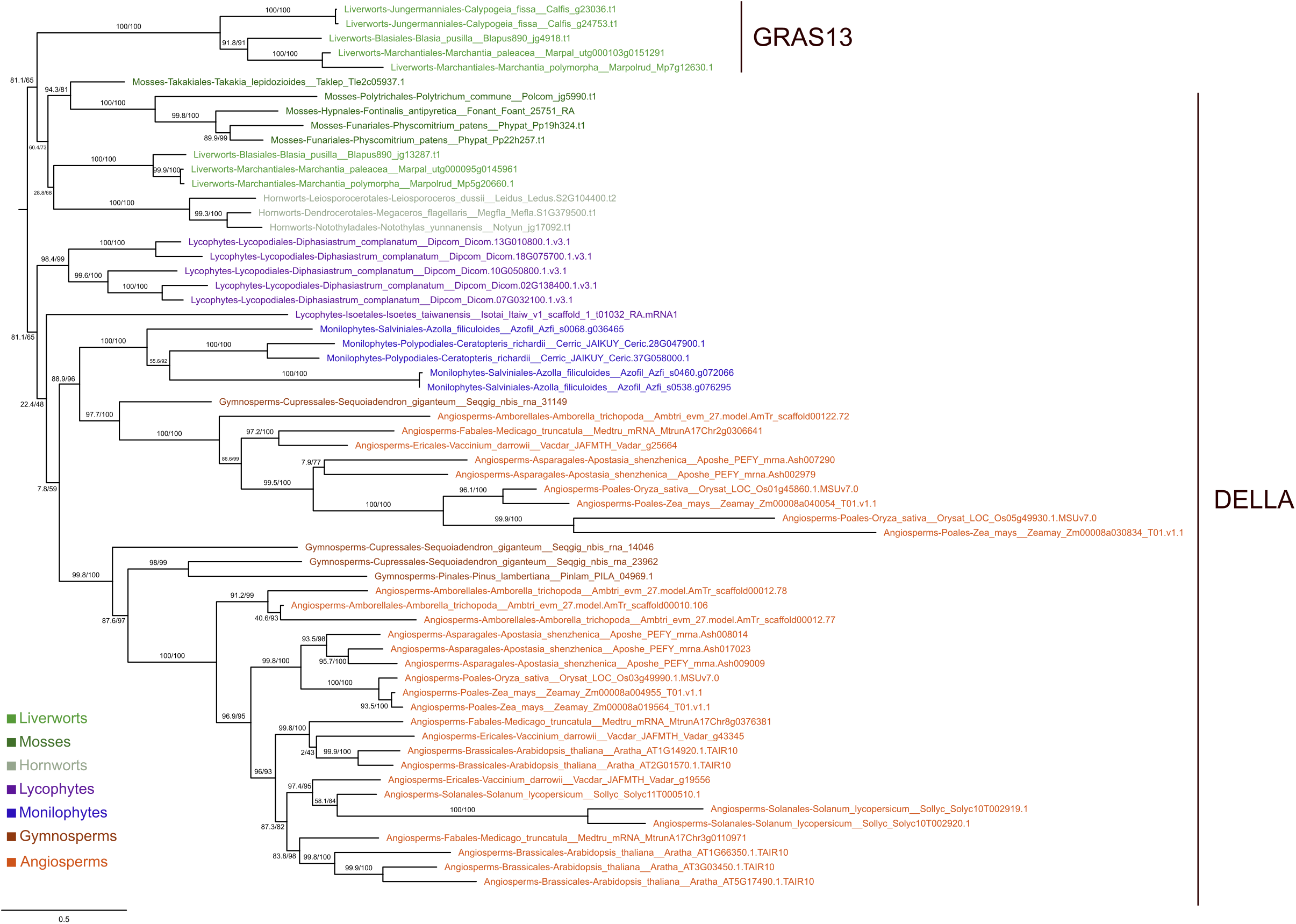
Phylogenetic tree of DELLA. Maximum-likelihood tree estimated from amino acid sequences of selected land plant species (model of substitution: JTT+F+I+G4, lnL: - 40062.661). Branch support values are shown next to branches (SH-like test/ultrafast bootstrap).

